# DrugMechDB: A Curated Database of Drug Mechanisms

**DOI:** 10.1101/2023.05.01.538993

**Authors:** Adriana Carolina Gonzalez-Cavazos, Anna Tanska, Michael D. Mayers, Denise Carvalho-Silva, Brindha Sridharan, Patrik A. Rewers, Umasri Sankarlal, Lakshmanan Jagannathan, Andrew I. Su

## Abstract

Computational drug repositioning methods have emerged as an attractive and effective solution to find new candidates for existing therapies, reducing the time and cost of drug development. Repositioning methods based on biomedical knowledge graphs typically offer useful supporting biological evidence. This evidence is based on reasoning chains or subgraphs that connect a drug to disease predictions. However, there are no databases of drug mechanisms that can be used to train and evaluate such methods. Here, we introduce the Drug Mechanism Database (DrugMechDB), a manually curated database that describes drug mechanisms as paths through a knowledge graph. DrugMechDB integrates a diverse range of authoritative free-text resources to describe 4,583 drug indications with 32,249 relationships, representing 14 major biological scales. DrugMechDB can be employed as a benchmark dataset for assessing computational drug repurposing models or as a valuable resource for training such models.

## 1 Introduction

Drug repositioning, the identification of novel uses of existing therapies, has become an increasingly attractive strategy to accelerate drug development Pushpakom et al. [2019]. By leveraging available genomics and biomedical domains, computational drug repositioning models have emerged as an unprecedented opportunity to analyze large amounts of data, reducing the time and effort required to identify repositioning candidates.

Computational repositioning models frequently rely on drug-drug and or disease-disease similarity Li et al. [2016], Li and Lu [2012]. However, the complex and contextual biological associations that underlie the relationship between a drug and a disease often require a more sophisticated explanation. To address this, biomedical knowledge graphs have emerged as a powerful tool capable of capturing biological associations that provide a more comprehensive understanding of the link between a drug and a disease Nicholson and Greene [2020].

Biomedical knowledge graphs consist of nodes representing biological concepts (such as genes, drugs, diseases, and pathways) and edges describing their relationship (such as drugs treating diseases, or diseases being associated with genes) Nicholson and Greene [2020]. Repositioning methods based on knowledge graphs leverage the biological associations captured on the network to provide supporting evidence for the model prediction. This is typically achieved by identifying subsets of reasoning chains or subgraphs within the larger network, providing a mechanistic rationale for why a particular drug might be effective against a particular disease, despite the absence of pre-existing evidence to validate the association Himmelstein et al. [2017].

However, one major challenge in determining the plausibility of the supporting evidence provided by biomedical knowledge graphs is the absence of a gold standard, well-defined collection of drug mechanisms. Such a reference point is necessary to evaluate the mechanistic accuracy of predictions made by repositioning models. While validation by domain experts is an alternative approach, it is a laborious and resource-intensive process that demands significant expertise.

Current efforts to construct biomedical networks integrate diverse knowledge bases Himmelstein et al. [2017], Santos et al. [2022], Yu et al. [2019], Zhu et al. [2020] or extract knowledge from literature using natural language processing (NLP) techniques Ernst et al. [2015], Percha and Altman [2018], Yuan et al. [2020]. However, there are several challenges in creating an accurate and comprehensive knowledge graph that serves as a high-quality benchmark dataset for repositioning discoveries. They often lack contextual information, not providing enough information about the relationship between a drug and a disease. Moreover, semantic interoperability is not present in high-quality, where concepts and terminologies within the network are unclear.

To fill this gap, we created Drug Mechanism Database (DrugMechDB), a manually curated database of drug mechanisms expressed as paths through a biomedical knowledge graph. The main focus of creating a detailed graph representation of mechanisms of action is to provide a useful community reference as a training or evaluation set for machine learning drug repurposing models. In this work, we present our first complete version of DrugMechDB capturing mechanisms for 4,583 drug-disease pairs. We provide a detailed analysis of the captured concepts and relationships within the paths of each record, elucidating the expressiveness of DrugMechDB. Lastly, we describe the utility of this database as a high-quality resource for repositioning methods based on knowledge graphs.

## 2 Methods

### 2.1 Indications composition

Indications were selected from DrugCentral (as downloaded on 2020-09-18)Ursu et al. [2016]. Each indication in DrugMechDB is represented as a directed graph, consisting of concepts and relationships (nodes and edges) that connect a drug to a disease that it treats. Each record contains several keys to produce a graph that can be accessed with any programming language (Figure 1).

**Figure 1:**
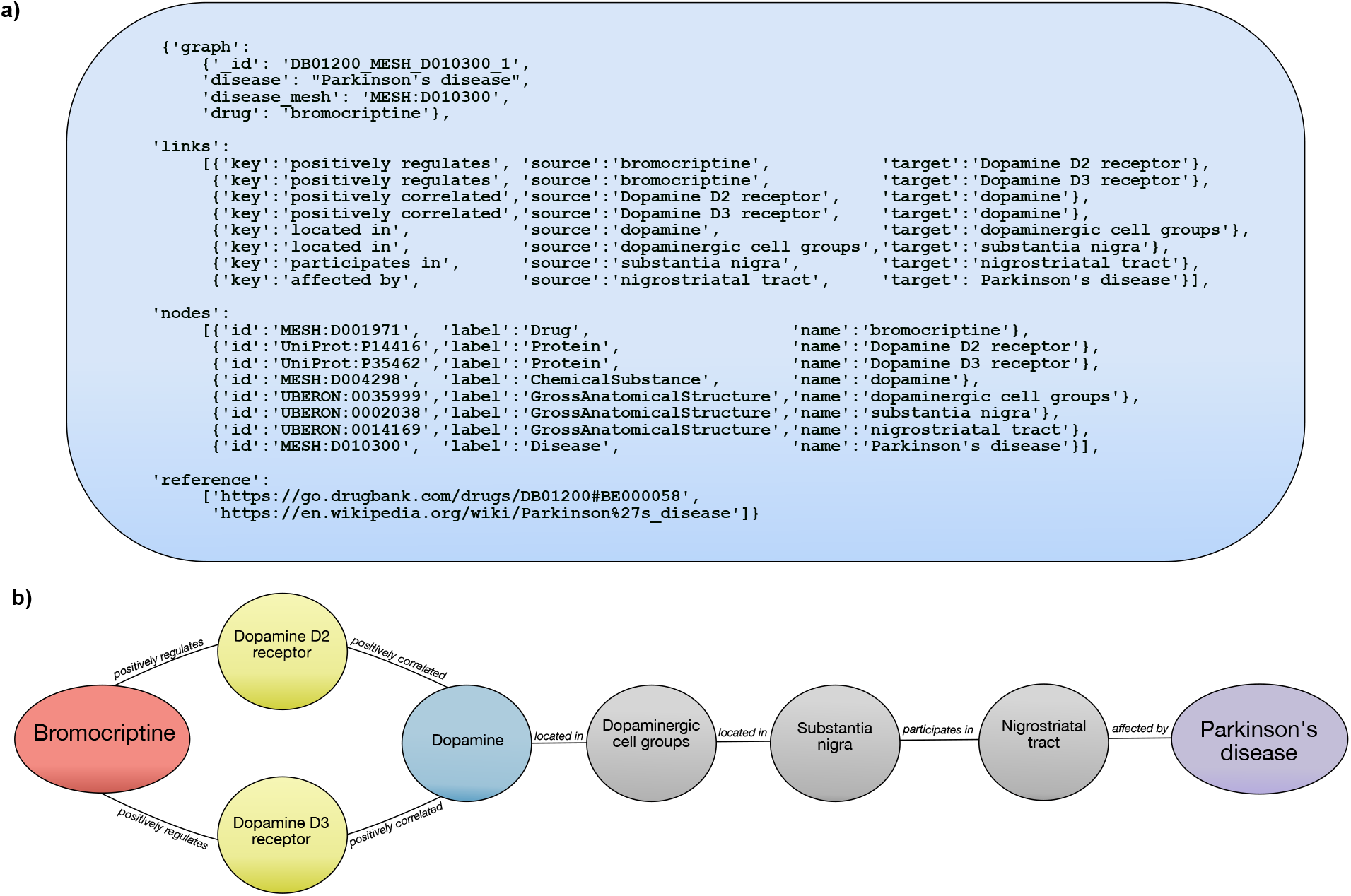
DrugMechDB indication components. A) Indication JSON formatting. Each record contains several keys that produce a graph that can be programmatically accessed: graph, links, nodes, and reference. B) Visualized example of one entry in DrugMechDB: a branching path from Bromocriptine to Parkinson’s disease.

### 2.2 Data Model

DrugMechDB provides researchers a consistent and structured information source on drug mechanisms. To achieve this goal, we adopted the Biolink Model (version 1.3.0) - a standardized hierarchy of biomedical entity classes that serves as a universal framework for biomedical data representation and linkage Unni et al. [2022]. The Biolink Model encompasses a wide range of entity types such as genes, proteins, diseases, drugs, and biological processes, and defines the predicates that describe the relationships between these entity types.

The standardization of data in DrugMechDB to the Biolink Model enables the mapping of concepts and relationships to a common vocabulary, thus allowing interoperability between various data sources. Consequently, researchers can easily combine data from DrugMechDB with other biomedical data sources that also employ the same data model, enabling researchers to perform comprehensive analyses and gain new insights into drug mechanisms of action. A list of the DrugMechDB concepts and corresponding relationships is found in Table 1.

**Table 1:** DrugMechDB concept types

| Node types | Abbreviation | Identifier Sources | Unique edge-types | Total edge count |
| --- | --- | --- | --- | --- |
| GrossAnatomicalStructure | A | Uber-anatomy ontology (UBERON)Mungall et al. [2012] | 24 | 534 |
| BiologicalProcess | BP | Gene Ontology (GO)Ashburner et al. [2000], gen [2021] | 38 | 8,235 |
| Cell | C | Cell Ontology (CL)Diehl et al. [2016] | 19 | 186 |
| CellularComponent | CC | Gene Ontology (GO)Ashburner et al. [2000], gen [2021] | 15 | 456 |
| Disease | D | Medical Subject Headings (MeSH)NLM [2021] | 12 | 147 |
| ChemicalSubstance | CS | Medical Subject Headings (MeSH)NLM [2021] ,<br>Chemical Entities of Biological Interest (ChEBI)Hastings et al. [2016] | 35 | 2,474 |
| Drug | DX | Medical Subject Headings (MeSH)NLM [2021] ,<br>DrugBankWishart et al. [2018] | 38 | 6,886 |
| GeneFamily | G | InterProPaysan-Lafosse et al. [2023], PfamMistry et al. [2021], | 21 | 958 |
| MolecularActivity | M | Gene Ontology (GO)Ashburner et al. [2000], gen [2021] | 21 | 474 |
| MacromolecularComplex | MC | Protein Ontology (PR)Natale et al. [2017] | 1 | 5 |
| Protein | P | UniProtCoudert et al. [2023] | 33 | 8,704 |
| PhenotypicFeature | PF | Human Phenotype Ontology (HP)Robinson et al. [2008] | 17 | 1,499 |
| Pathway | PW | Reactome Pathway (REACT)Jassal et al. [2020] | 20 | 348 |
| OrganismTaxon | T | NCBITaxonSchoch et al. [2020] | 5 | 1,343 |
|  |  |  | Total | 32,249 |

### 2.3 Path curation

One of the most important aspects of the curation process is relying on sources that have undergone rigorous review and validation processes, ensuring that the information is accurate, reliable, and up-to-date. The curation of DrugMechDB was based on a diverse range of authoritative free-text resources, including review articles, DrugBank Wishart et al. [2018], GeneOntology Ashburner et al. [2000], UniProt Coudert et al. [2023], Reactome Jassal et al. [2020], and well-sourced Wikipedia articles Vrandečić [2012]. Primary literature sources containing experimental results were excluded, ensuring that only highly curated and high-confidence information was included. This exclusion helped to preserve the quality and accuracy of the information, prioritizing quality over quantity.

The process of defining relationships that describe a drug’s action from free-text descriptions can be subjective, which can result in inconsistent annotations. To ensure uniformity in path representations among DrugMechDB entries, we established a formal curation guide. Briefly, we ensured to keep the order of interactions to reflect cause and effect between concepts, extraneous interactions or information was removed, and multiple related concepts were summarized in a single all-encompassing concept. A guideline is provided within the code repository at Github (/SuLab/DrugMechDB/blob/main/CurationGuide).

In any manual process, the possibility of errors always exists. While the curation process of DrugMechDB is very detail-oriented and careful, we also validated all node identifier names against an authoritative resource to increase standardization and reduce human error. We used the node normalization service (version 2.0.9) from the Biomedical Data Translator program as a primary source to validate node identifier namesFecho et al. [2022]; more details are provided within the code repository at Github (/TranslatorSRI/NodeNormalization).

## 3 Data Records

The first completed DrugMechDB version (2.0.0) is available at Github (https://github.com/SuLab/DrugMechDB) and Zenodo (DOI 10.5281/zenodo.7868996). Curated indications are found in file named indication_paths.json The structure of the provided JSON file is shown in Figure 1a.

### 3.1 Concepts and relationships captured in DrugMechDB indications

DrugMechDB is a large curated biomedical network capturing 4,583 unique indications between 1,580 drugs and 744 diseases. DrugMechDB contains 32,588 nodes of 14 biological concept types, and 32,249 edges classified into 71 different edge types. We provide a breakdown of the number of edges by concept type in Table 1.

Out of the 14 concept types in DrugMechDB, the BiologicalProcess concept type appears most frequently as a node on the graph, comprising 24.55% of the total nodes. The Protein concept type comes in a close second, representing 21.53% of the total nodes (Figure 2a). These results highlight the essential role of proteins and biological processes in drug mechanisms of action. Proteins act as the primary molecular effectors of biological processes, which are fundamental building blocks of cellular functions.

**Figure 2:**
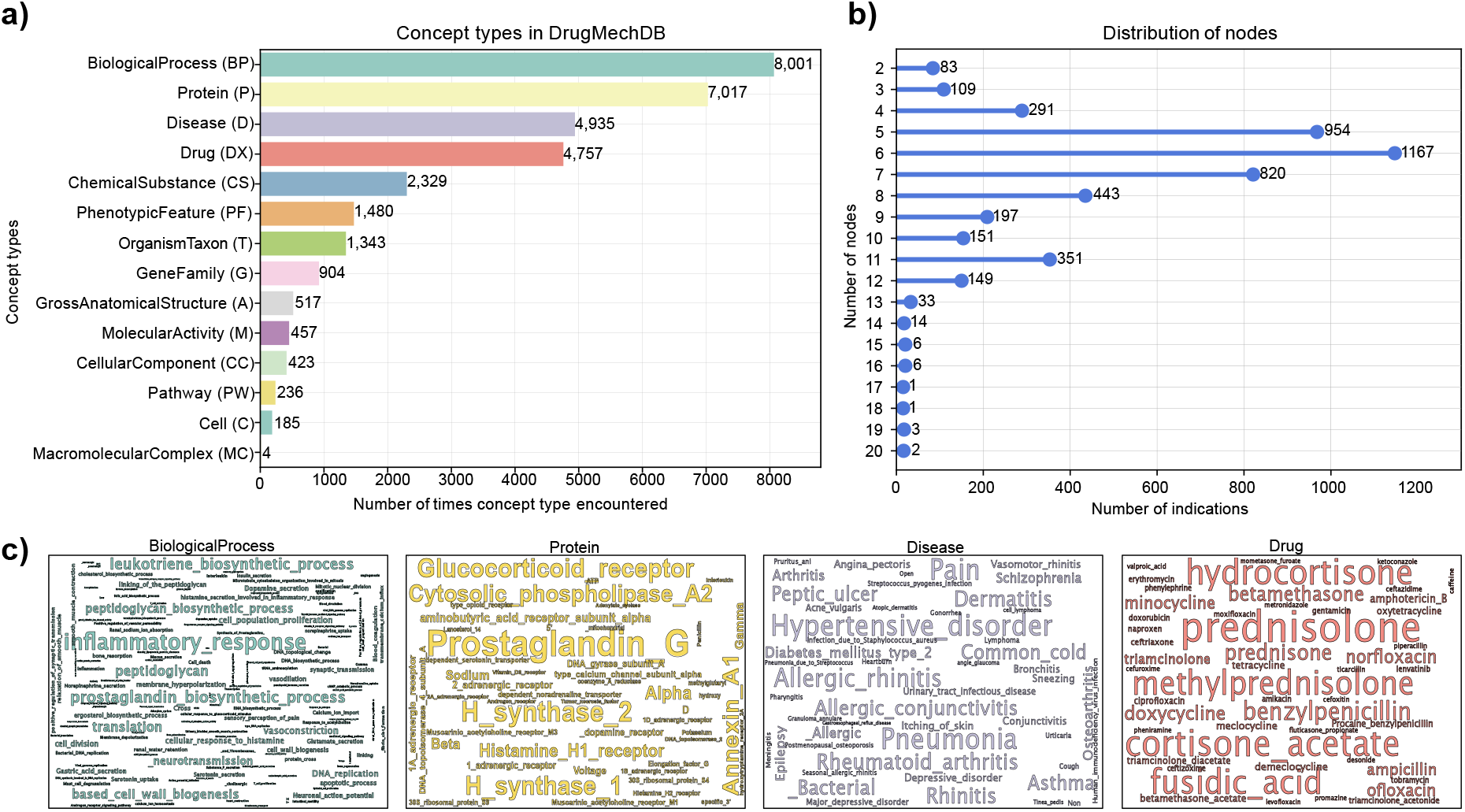
Concept types in DrugMechDB. A) Total number of unique nodes found by concept type within DrugMechDB. Abbreviations are shown in parentheses. B) Distribution of encountered nodes per indications. C) Most frequent nodes within BiologicalProcess (blue), Protein (yellow), Disease (purple), and Drug (red) universe

Among the total 129 pairing of concept types in DrugMechDB, the most common connection occurs between a Protein to a BiologicalProcess concept (e.g., c-Kit protein connected to cellular proliferation biological process), with 4,695 examples, accounting for 14.55% of the total associations (Figure 3a). Additionally, out of the total 725 meta-edges, the most frequent type occurs connecting a Protein to a BiologicalProcess concept through a positively regulates edge-type (e.g., c-Kit positively regulates cellular proliferation), representing 11.29% of the total meta-edges in DrugMechDB (Figure 3b).

**Figure 3:**
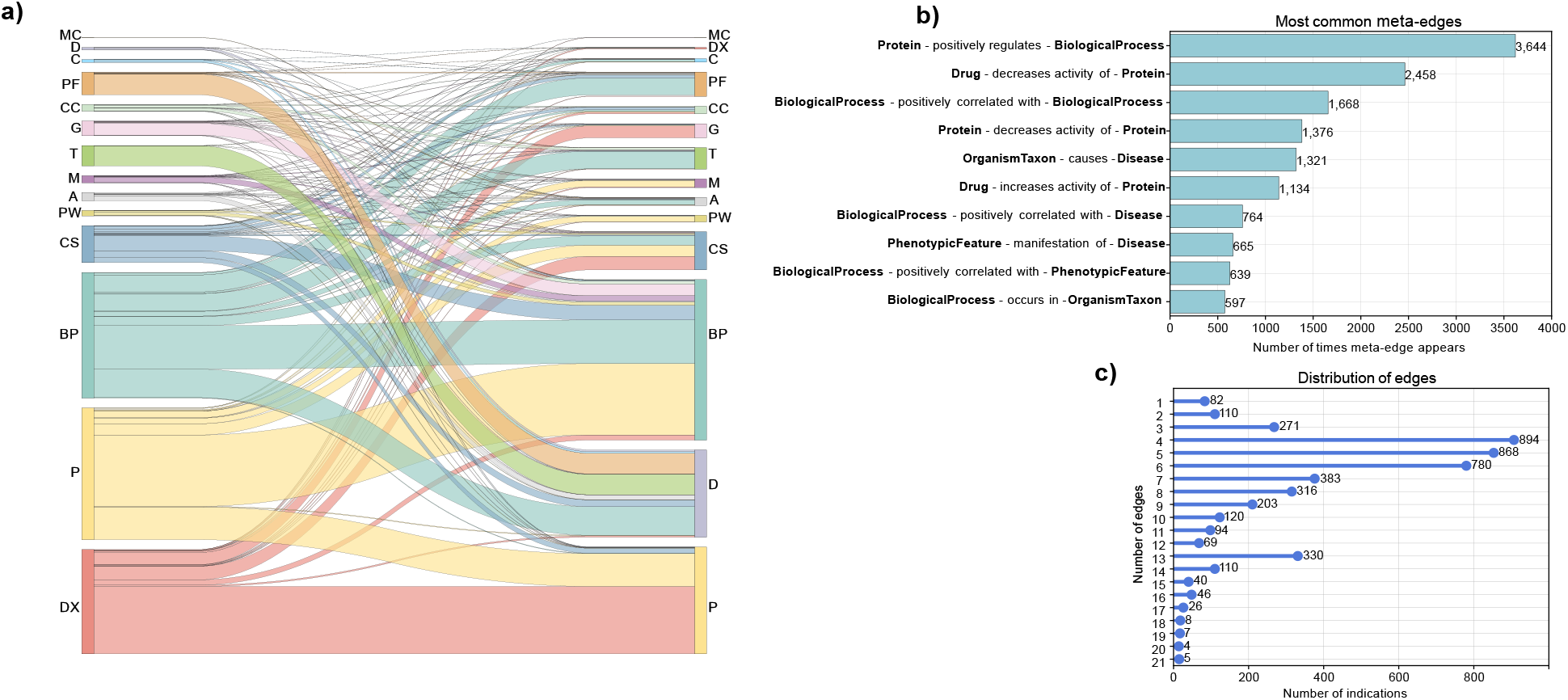
Edge types in DrugMechDB. A) Sankey diagram showing existing edges between all concept types, thickness equates to the number of connections. Concept types are abbreviated. B) Top ten encountered meta-edges. C) Distribution of edges per indications.

The complexity of interactions involved in drug-disease associations can lead to a wide variation in the number of nodes and edges. To illustrate this variation, Figure 2b and Figure 3c depict the distribution of the number of nodes and edges captured in DrugMechDB indications, respectively. As shown, some indications are relatively simple, with only a few nodes and edges, while others are much more complex, with many interconnected nodes and edges, reflecting the complexity of the biological connections.

### 3.2 Mechanistic paths captured in DrugMechDB indications

We conducted an evaluation of the 5,666 mechanistic paths explaining curated drug actions. In DrugMechDB, mechanistic paths are categorized into 297 types based on the sequence of concept types, disregarding the edges between them. We identified that the mechanistic path Drug-Protein-BiologicalProcess-Disease is the most commonly occurring sequence of concept types among indications, representing 12.27% of the total DrugMechDB paths (Figure 4a). As depicted above, Protein and BiologicalProcess concept types are the two most commonly occurring node types on DrugMechDB, emphasizing their crucial function in drug mechanisms of action. For instance, Figure 5a depicts a visualized example of one DrugMechDB entry for this mechanistic path type.

**Figure 4:**
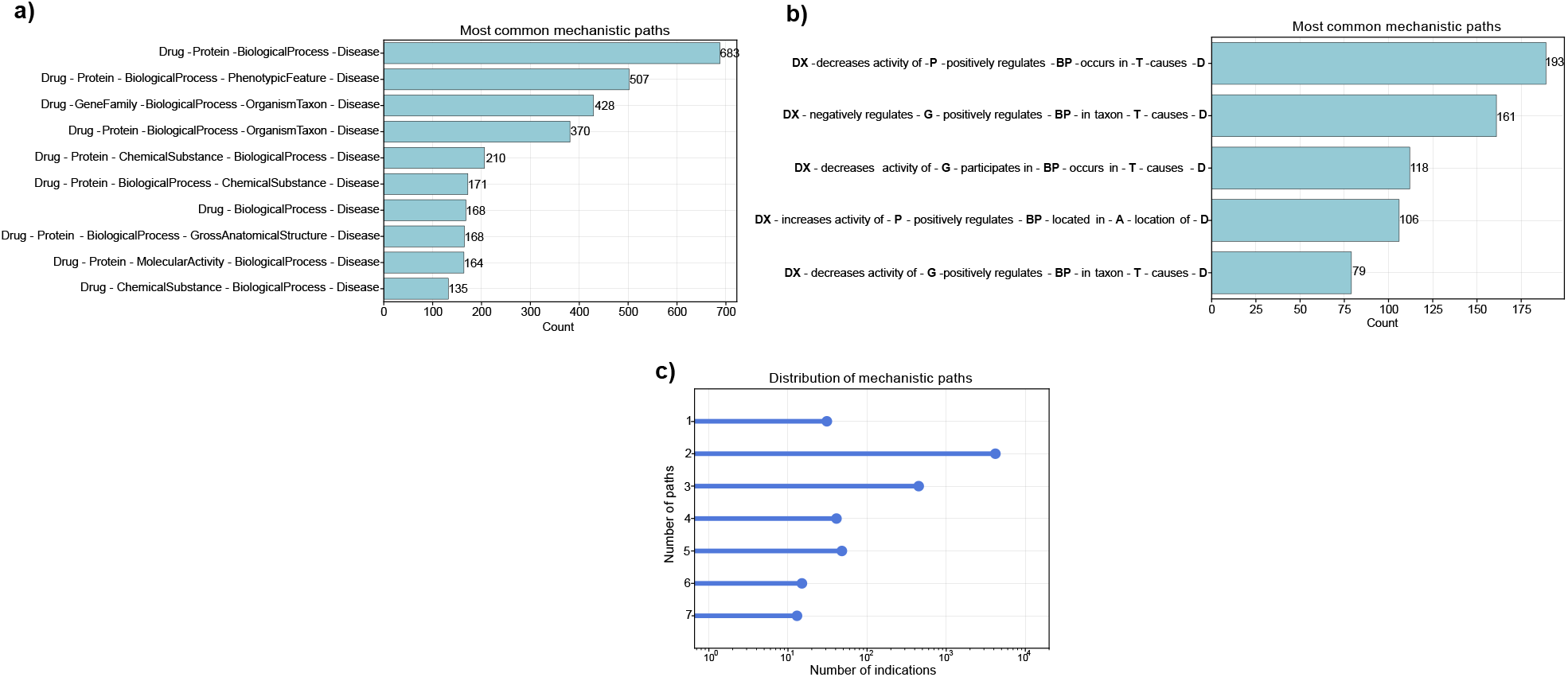
Mechanistic paths in DrugMechDB. A) Top ten most commonly occurring mechanistic paths, determined solely by the sequence of concept types. B) Top ten most commonly occurring mechanistic paths, determined considering edges connecting concept types. Concept types are abbreviated. C) Distribution of captured paths per indications.

**Figure 5:**
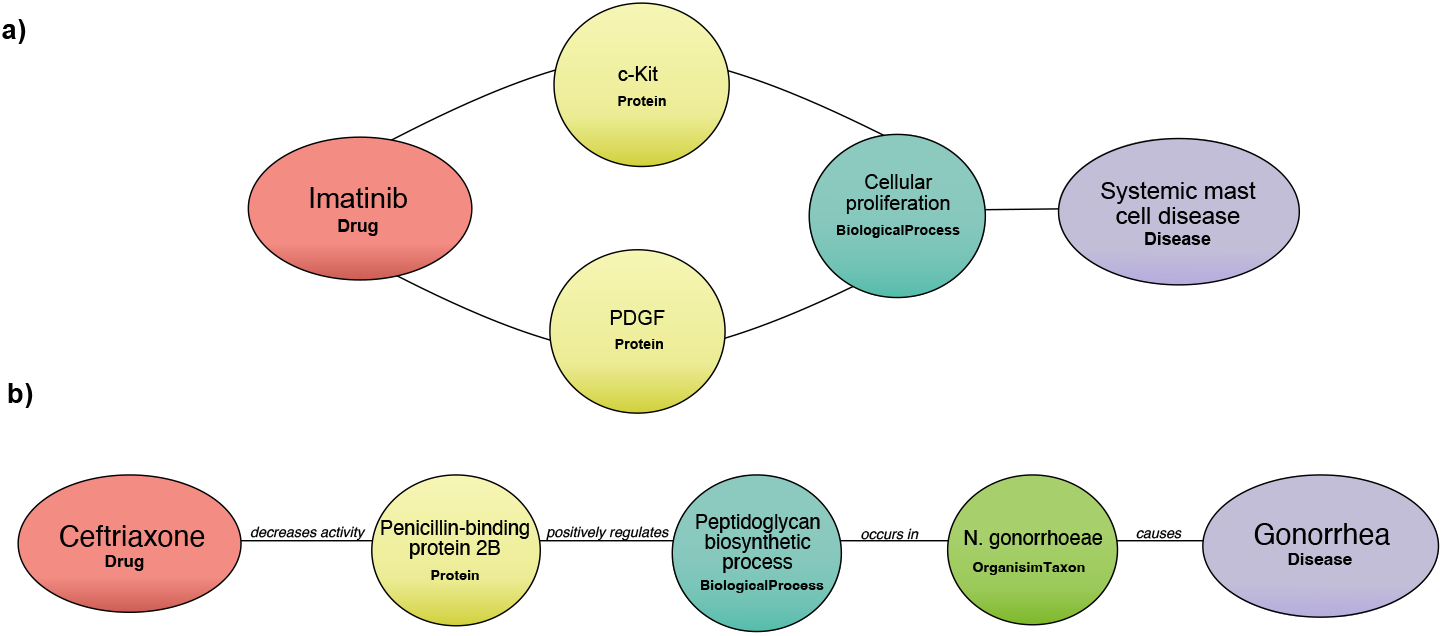
Mechanistic paths in DrugMechDB. A) An example of the most common occurring sequence of concept types: a path from Imatinib to Systemic mast cell disease. B) An example of the most common occurring sequence of concept types and edge types: a path from Ceftriaxone to Gonorrhea.

Moreover, in DrugMechDB, paths are further classified into 1,457 unique mechanistic types based on the edge type connecting concept types. We identified that the mechanistic path Drug -decreases activity of-Protein -positively regulates-BiologicalProcess -occurs in-OrganismTaxon -causes-Disease is the most commonly occurring type, accounting for 3.46% of all paths (Figure 4b). For instance, Figure 5b depicts a visualized example of one DrugMechDB entry for this mechanistic path type.

It is worth mentioning that some drugs achieve their therapeutic benefits by engaging in multiple simultaneous interactions. This might involve blocking multiple targets that work together to produce their effect or affecting multiple unrelated pathways that would have little effect on the disease if targeted separately. DrugMechDB represents this type of scenario through a branching path, with an average of two branching paths outlining the link between a drug and a disease (Figure 4c).

## 4 Technical Validation

### 4.1 Systematic validation of DrugMechDB associations

Validating the reliability of a knowledge graph is a crucial step that ensures the correctness of the captured information. In this work, we assessed the accuracy of captured DrugMechDB associations by comparing them to existing data sources. For this, we leverage an external biomedical knowledge graph: Mechanistic Repositioning Network (MechRepoNet) Mayers et al. [2022].

Briefly, MechRepoNet is a comprehensive biomedical knowledge graph that was constructed by integrating 18 different data sources and using Biolink Model for standardization. Given that MechRepoNet encompasses a wider network that spans various domains, we employed it as an external benchmark for verifying the plausibility of the associations recorded in DrugMechDB.

Evaluating association types between concept types (ignoring edge predicates), we found that 2,924 (28.71%) of the 10,184 unique associations captured in DrugMechDB are also contained within MechRepoNet. To demonstrate that DrugMechDB associations are broadly consistent with the knowledge captured in MechRepoNet, we conducted a bootstrapping analysis. For each DrugMechDB association type, nonparametric bootstrapping was applied to sample simulated association types (with replacement) to calculate the percentage of matching with MechRepoNet. This procedure was repeated 1,000 times to construct a percentage distribution from which the mean and 99% CI were calculated. The p-value was calculated as the fraction of the distribution in which the simulated percentage of matching was greater than or equal to the observed percentage. Results in Table 2 show that the average p-value of the ten most frequent association types is less than 0.001, demonstrating that observed overlapping between DurgMechDB and the broader knowledge captured by MechRepoNet is unlikely to occur by chance.

**Table 2:** Validation of the ten most frequent DugMechDB association types

| DrugMechDB association type | DMDB count | MechRepoNet overlap (%) | Mean bootstrapping overlap (99(%) CI) | P-value |
| --- | --- | --- | --- | --- |
| Protein-BiologicalProcess | 4,699 | 40.15 | 2.21 (1.78-2.66) | < 0.001 |
| Drug-Protein | 4,475 | 60.78 | 4.74 (4.02-5.45) | < 0.001 |
| BiologicalProcess-BiologicalProcess | 2,889 | 0.38 | 0.29 (0.10-0.55) | 0.137 |
| Protein-Protein | 2,166 | 2.15 | 0.09 (0-0.13) | < 0.001 |
| BiologicalProcess-Disease | 1,897 | 56.48 | 39.40 (36.90-41.96) | < 0.001 |
| PhenotypicFeature-Disease | 1,352 | 6.41 | 0.002 (0-0.07) | < 0.001 |
| OrganismTaxon-Disease | 1,340 | 30.76 | 1.37 (0.82-1.94) | < 0.001 |
| BiologicalProcess-OrganismTaxon | 1,161 | 11.59 | 6.08 (4.9-7.40) | < 0.001 |
| BiologicalProcess-PhenotypicFeature | 1,136 | 23.59 | 18.71(16.37-21.12) | < 0.001 |
| ChemicalSubstance-BiologicalProcess | 972 | 9.13 | 3.46 (2.36-4.62) | < 0.001 |

The association type BiologicalProcess-BiologicalProcess has the least overlap among the most frequent DrugMechDB association types, highlighting that MechRepoNet does not cover all curated association types of DrugMechDB. To incorporate the missing information in MechRepoNet, we propose using DrugMechDB as a roadmap, helping to prioritize the most significant relationships involved in drug mechanisms and facilitating the integration of biomedical sources.

In summary, DrugMechDB is a comprehensive resource that provides human interpretable explanations when producing computational repurposing predictions, it has the potential to help domain experts to better assess whether a model’s candidate provides enough biological evidence. We believe that DrugMechDB offers several advantages. First, it serves as a useful resource for researchers looking to understand drug pharmacodynamics. Second, it is a valuable training data set that can be incorporated into drug repositioning models that focus on providing supporting plausible reasoning chains. Lastly and as described above, DrugMechDB functions as a roadmap for knowledge graph expansion, helping to prioritize biological associations that most commonly appear in curated drug mechanisms.

## 5 Code availability

A web interface to DrugMechDB can be found at https://sulab.github.io/DrugMechDB/. The code to reproduce results, together with documentation is available at https://github.com/SuLab/DrugMechDB. In addition, contributions of curated mechanistic paths can be done by pull request to the file submission.yaml. Additional submission details are available at SuLab/DrugMechDB/blob/main/SubmissionGuide.md

## Acknowledgements

This work was supported by funding from the National Center for Advancing Translational Sciences (NCATS) under awards OT2TR003427 and UL1TR002550, and from the National Institutes of Aging (NIA) under award R01AG066750. The content is solely the responsibility of the authors and does not necessarily represent the official views of the National Institutes of Health.

## Author contributions statement

AT, MM, BS, DC-S, US and LJ retrieved and curated indications. AG-C, MM, and PR wrote analysis tools and performed the analysis. AG-C and AS wrote the manuscript. MM and AS conceptualized and designed the study. All authors have read and approved the manuscript.

